# Subthalamic stimulation evoked cortical responses relate to motor performance in Parkinson’s disease

**DOI:** 10.1101/2022.06.28.497955

**Authors:** Bahne H. Bahners, Rachel K. Spooner, Christian J. Hartmann, Alfons Schnitzler, Esther Florin

## Abstract

Subthalamic deep brain stimulation is effective in alleviating motor symptoms in Parkinson’s disease. Establishing the clinically best stimulation settings often requires time-consuming test sessions and creates a need for biomarkers to optimize this process. While stimulation-evoked cortical responses have been proposed as such a neurophysiological marker, their relationship to motor performance has not yet been studied systematically. For this aim, we recorded finger-tapping movements and cortical responses evoked by different stimulation amplitudes of 22 patients with Parkinson’s disease using magnetoencephalography. The motor cortex amplitude was a significant predictor of a higher finger tap frequency and a more regular tapping profile. In addition, subthalamic stimulation evoked responses in the inferior and middle frontal gyrus, and the supplementary motor area. While earlier studies relied on a limited cortical coverage, we reveal a cortical distribution of responses that aligns with the basal ganglia-thalamo-cortical network. Our study sheds light on the relationship between cortical responses evoked by subthalamic stimulation and motor performance based on objective quantitative parameters. Stimulation-evoked responses could guide clinical programming in the future.

## Introduction

Deep brain stimulation (DBS) of the subthalamic nucleus (STN) is an established therapy for Parkinson’s disease.^1^ However, determining the optimal stimulation setting can be a time-consuming trial-and-error process.^3^ Therefore, biomarkers that can guide the optimization of stimulation settings are highly desireable.^4^ STN beta band activity has been proposed as such a marker several years ago, due to its consistent positive correlation with symptom severity and negative correlation to clinical improvement through DBS.^5,6^ Still, chronic access to electrophysiological recordings from the basal ganglia is limited and only possible with novel sensing-enabled neurostimulators.^6^

Thus, there is a need to define non-invasive biomarkers, for instance through the analysis of DBS-evoked cortical responses.^7–10^ Previous studies indicate that responses with latencies of 2 to 10 ms – resulting from antidromic hyperdirect pathway activation – are higher for stimulation contacts that elicit a therapeutic effect.^11^ Similarly, in rodent models of Parkinson’s disease the amplitude of motor cortex evoked responses was associated with improved motor symptoms.^12,13^ Responses at longer latencies (>20ms) may represent an orthodromic synaptic transmission via the basal ganglia-thalamo-cortical loop.^11^ However, neither the direct relationship to objective measures of motor performance nor the precise cortical distribution of stimulation-evoked responses have been studied so far.

We hypothesized that i) stimulation-evoked responses of the motor cortex represent a marker of antidromic as well as orthodromic cortical activation and ii) that the amplitude of these responses relates to motor performance at high-frequency stimulation. Making use of the high temporal and spatial resolution of magnetoencephalography (MEG), we analyse cortical evoked responses and relate them to movement performance based on objective quantitative parameters extracted from accelerometer recordings in a cohort of patients with Parkinson’s disease.

## Materials and methods

### Patients

A group of 22 patients with Parkinson’s disease (19 male, 3 female, 64 ± 9 years) was recruited at the Center for Movement Disorders and Neuromodulation (University Hospital Düsseldorf). Mean disease duration was 12 ± 6 years, mean time after bilateral STN-DBS implantation was 26 ± 14 months. Motor impairment was assessed using the unified Parkinson’s disease rating scale (UPDRS III: Medication OFF/Stimulation OFF: 40 ± 11 *vs*. Medication ON/Stimulation OFF: 29 ± 13 *vs*. Medication ON/Stimulation ON: 15 ± 8). During MEG recordings, patients were in their best medication ON state (mean levodopa-equivalent daily dose: 661 ± 324mg). All patients gave their prior written informed consent, the study was approved by the local ethics committee (study number: 2019-629_2), and performed in accordance with the Declaration of Helsinki.^14^

### Behavioural experiment and accelerometer analysis

A triaxial accelerometer^15^ (ADXL335 iMEMS Accelerometer, Analog Devices Inc., Norwood, MA, USA) was attached to the tip of the patient’s right index finger. The patients were asked to tap 10 times their right index finger onto their thumb and to perform the movement as large, fast and regular as possible (item 3.4, UPDRS III). During tapping the left STN was stimulated with an omnidirectional monopolar montage on the patient’s clinically selected contact with 130 Hz and a pulse width of 60µs. The tapping sequence was performed and recorded for each tested stimulation amplitude (0.5, 1.0, 2.0, 3.0, and 4.0 mA). At stimulation amplitudes that elicited immediate sustained side effects, the task was not performed and these conditions were omitted from further analysis.

The accelerometer data was epoched based on a video of the session, visually inspected for artefacts, and processed using custom-written MATLAB scripts (2021a, MathWorks Inc.). To evaluate the tap variability within each block we used a fixed-threshold algorithm, with which the onset of single finger taps was detected based on the 95% percentile across time of the variation in accelerometer vector length. Tap detection results were visually inspected. In case the algorithm missed taps, the percentile threshold was lowered accordingly. The time between two consecutive taps was determined (inter-tap interval) and each inter-tap interval was transformed into taps per second (1000ms/inter-tap interval). Based on the taps per second we determined the average tap frequency during a tapping block and as index of the tap variability the coefficient of variation over the tapping block (i.e., standard deviation in tap frequency/mean tap frequency x 100). Tap frequency and variability were chosen as movement metrics, because they reflect important criteria to rate motor performance of item 3.4 in the UPDRS.

### MEG acquisition and data analysis

After the accelerometer testing, the patient was seated in a whole-head MEG system with 306 sensors (Elekta Oy, Helsinki, Finland) and neuromagnetic activity was measured in short acquisition runs of 40 seconds per stimulation setting. We used a low-frequency (6 Hz) monopolar stimulation of the left STN to enable the analysis of stimulation-evoked responses at longer latencies.^7^ Except for the stimulation frequency, the same monopolar stimulation settings as used during the accelerometer testing were applied (i.e., stimulation amplitude, contact, pulse width). Due to movement-related artefacts in the right-sided sensors, resulting from the DBS hardware^16^, we focused our analysis on responses evoked by left STN stimulation and right-hand finger tapping.

MEG data was sampled at 5,000 Hz with a high-pass of 0.1 Hz and a low-pass of 1,660 Hz. Prior to the MEG recording, patients’ head shape and head position indicator coils were digitized with a 3D-digitizer (Fastrak Digitizer, Polhemus, Vermont, USA). Eye movements (EOG) and heart activity (ECG) were monitored throughout the measurements. MEG data analysis was performed with Brainstorm.^17^ MEG recordings from gradiometers (204) and magnetometers (102) were inspected for channel jumps and subjected to temporal signal space separation (tSSS) using MNE python’s version in Brainstorm.^18^ A notch filter was applied to reduce power line noise at 50 Hz and its harmonics up to 300 Hz. Signal space projection was used to eliminate cardiac and blink artefacts.^19^ An additional EMG channel placed above the pulse generator was used for the detection of stimulation pulses. Trials were defined as a period of 100 ms after the stimulation pulse with a baseline from -50 ms to -5 ms. For further analysis on average 228 ± 31 clean trials per subject and condition were included. The number of trials did not differ significantly between stimulation amplitudes (0.5mA: 227±20 trials, 1mA: 233±15 trials, 2mA: 225±27 trials, 3mA: 229±34 trials, 4mA: 226±49 trials). Evoked responses for each stimulation amplitude and MEG sensor were obtained by averaging across the corresponding trials.

The significant time-windows of interest used for subsequent source analyses were determined through a two-stage, data-driven approach.^20^ First, for each sensor a paired-sample t-test against baseline was computed on the evoked responses pooled across all patients and tested stimulation amplitudes. Second, cluster-based permutation testing was employed to control for multiple comparisons (initial threshold: *P*< 0.05, permutations: 1,000, minimum duration: 5 ms).^21^ After the DBS pulse, activity significantly changed compared to baseline in three time-periods in left-sided sensors ipsilateral to the stimulation: 2.4 to 9.8 ms, 10.0 to 17.6 ms, and 18.6 to 27.8 ms. Due to the remnants of the stimulation artefact, we changed the beginning of the first time-window to 3.6 ms (Figure 1 A, B, F, G). The maximum t-value across time and sensors within the first window was 5.6 ms, 11.8 ms for the second and 21.8 ms for the third (from now on referred to as M5, M10 and M20, respectively).

**Figure 1.**
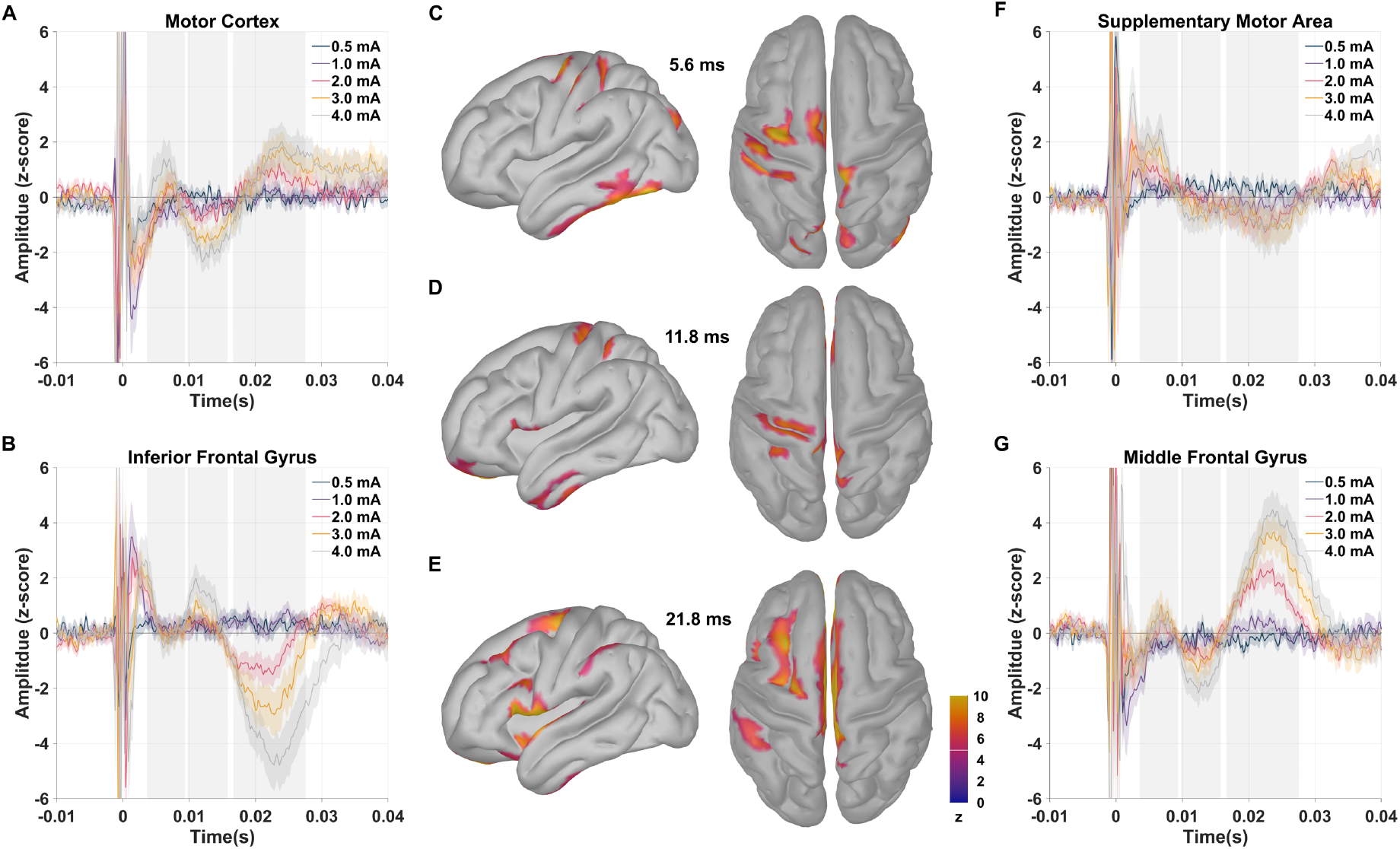
Grand average source time series and cortical pattern of stimulation-evoked responses. (**A, B, F, G**) Source time series across patients and standard errors of the mean at the five different stimulation amplitudes tested. The grey-shaded areas indicate the time windows from which the individual peak amplitudes were extracted based on sensor-level analyses. Time series were extracted from four regions of interest. (**C-E**) Source images for the evoked response peak latencies at 5.6 (**C**), 11.8 (**D**) and 21.8 ms (**E**). The absolute z-scored amplitudes are depicted on a MNI template (ICBM125 2009c Nonlinear Asymmetric) using a threshold of 5.0 (z-score) for visualisation purposes as indicated by the white line in the colour bar.

Cortical surfaces were estimated from individual MRIs using CAT12,^22^ and co-registered with the MEG recordings using an iterative closest-point rigid-body registration in Brainstorm. Next, the forward problem was solved with an overlapping spheres head model and 15,000 cortical sources were reconstructed with a linearly constrained minimum variance (LCMV) beamformer from the evoked responses on the sensor level for each patient and each stimulation amplitude (22 × 5 source maps).^23^ All individual source level data were projected to MNI space (ICBM125 2009c Nonlinear Asymmetric)^24^ using Shepard’s method as implemented in Brainstorm. We created a grand-average source map across all patients and stimulation amplitudes and z-score baseline normalized it. We determined the vertex with the maximum value for the peak latencies of the M5, M10 and M20 response on the hemisphere ipsilateral to the stimulation (left) within the following regions: motor cortex, supplementary motor area, middle frontal gyrus, and inferior frontal gyrus. These regions of interest were selected based on the grand-average source map at peak latencies of M5, M10 and M20 with a z-score larger than 6 (Figure 1 C-E). From all of the source maps (each patient and each stimulation amplitude) we then extracted and averaged the 30 vertex time series around the maximum in each region. These mean source time series were baseline normalized using a z-score baseline normalization ([-50ms, -5ms]). Afterwards, we used the absolute extracted time series to determine the evoked response amplitude maxima in each patient and stimulation condition within the previously defined time-windows.

### Statistics

For statistical analyses we used MATLAB (2021a, MathWorks Inc.). We estimated linear mixed effects models of movement outcomes (e.g., tap frequency, variability) as a function of evoked cortical response amplitude (continuous variable), DBS stimulation amplitude (continuous variable), and their interaction, with subject included as a random effect. Of note, separate models were run for each response outcome measure, cortical region, and latency. *P*-values were corrected for multiple comparisons across these dimensions using false discovery rate correction^25^ and only the adjusted *P*-values are reported in the following.

### Data availability

The anonymized raw data are available on request and in accordance with data privacy statements signed by all patients.

## Results

The DBS pulse consistently elicited three ipsilateral cortical responses peaking at 5.6 ms (M5), 11.8 ms (M10), and 21.8 ms (M20), see Figure 1 C-E. The cortical activation of M5 involved the motor cortex and the supplementary motor area (Figure 1 C). The M10 was mainly located in the motor cortex, and the M20 included the supplementary motor area as well as the middle and inferior frontal gyrus (Figure 1 E).

Next, we evaluated the relation between cortical evoked responses and DBS stimulation amplitude on behaviour using linear mixed effects models. Both the M5 and M20 response in the motor cortex and the supplementary motor area were significant predictors of tap variability (Figure 2 A, C, D), such that greater M5 and M20 responses predicted greater consistency in finger tapping frequency (M5: motor cortex: *b* = -8.60 ±3.12, *P* = 0.043; supplementary motor area: *b* = -8.60 ±2.99, *P* = 0.043; M20: motor cortex: *b* = -16.10 ±4.13, *P* = 0.005; supplementary motor area: *b* = -8.59 ±3.16, *P* = 0.043). The motor cortex M10 response was a significant predictor of the tap frequency (Figure 2 B), with greater evoked responses related to greater tap frequencies (*b* = 0.23 ±0.09, *P* = 0.043). In addition, there was a significant interaction of stimulation amplitude with evoked response in case of the tap variability (*b*= 4.60 (±1.37), *P =* 0.028).

**Figure 2.**
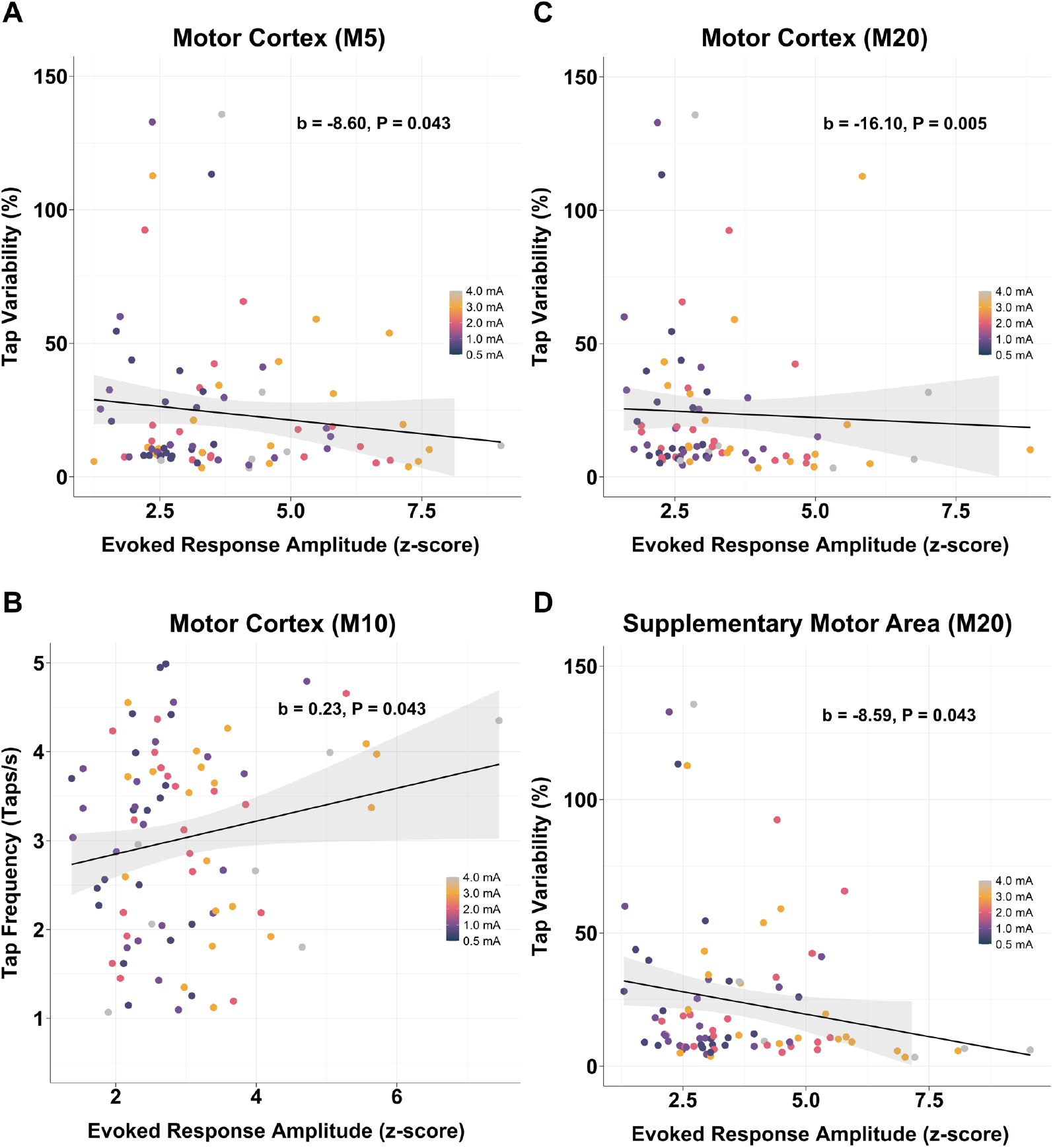
Response amplitudes relate to motor performance. Scatter plots depicting finger tap variability (**A, C, D**) and tap frequency (**B**) as a function of cortical M5 (**A**), M10 (**B**) and M20 (**C, D**) response amplitudes in the motor cortex (**A-C**) and the supplementary motor area (**D**) with respective regression lines and 90%-confidence intervals. The colour bar indicates the stimulation amplitude. In each panel the coefficients *b* from the linear mixed effects models and the respective *P*-values are provided.

**Table 1.** Estimated coefficients from linear mixed effects models.

| Region | Stimulation Amplitude | Response Amplitude | Stimulation*Response |
| --- | --- | --- | --- |
| **Tap frequency (tap/s): M5 response** |  |  |  |
| MI | 0.09 ($\pm 0.08$), $P = 0.412$ | 0.06 ($\pm 0.06$), $P = 0.397$ | -0.01 ($\pm 0.02$), $P = 0.527$ |
| SMA | 0.10 ($\pm 0.07$), $P = 0.322$ | 0.05 ($\pm 0.06$), $P = 0.449$ | -0.01 ($\pm 0.02$), $P = 0.506$ |
| MFG | 0.12 ($\pm 0.08$), $P = 0.309$ | 0.05 ($\pm 0.05$), $P = 0.401$ | -0.02 ($\pm 0.02$), $P = 0.490$ |
| IFG | -0.02 ($\pm 0.08$), $P = 0.882$ | -0.06 ($\pm 0.06$), $P = 0.401$ | 0.03 ($\pm 0.03$), $P = 0.463$ |
| **Tap frequency (taps/s): M10 response** |  |  |  |
| MI | 0.16 ($\pm 0.08$), $P = 0.256$ | 0.23 ($\pm 0.09$), **$P = 0.043$** | -0.05 ($\pm 0.03$), $P = 0.187$ |
| SMA | -0.01 ($\pm 0.10$), $P = 0.932$ | -0.07 ($\pm 0.08$), $P = 0.440$ | 0.03 ($\pm 0.03$), $P = 0.506$ |
| MFG | 0.13 ($\pm 0.08$), $P = 0.296$ | 0.08 ($\pm 0.08$), $P = 0.397$ | -0.02 ($\pm 0.03$), $P = 0.490$ |
| IFG | -0.08 ($\pm 0.08$), $P = 0.445$ | -0.13 ($\pm 0.06$), $P = 0.124$ | 0.05 ($\pm 0.02$), $P = 0.187$ |
| **Tap frequency (tap/s): M20 response** |  |  |  |
| MI | 0.15 ($\pm 0.08$), $P = 0.256$ | 0.13 ($\pm 0.08$), $P = 0.201$ | -0.04 ($\pm 0.03$), $P = 0.325$ |
| SMA | 0.11 ($\pm 0.07$), $P = 0.305$ | 0.05 ($\pm 0.06$), $P = 0.440$ | -0.02 ($\pm 0.02$), $P = 0.490$ |
| MFG | 0.15 ($\pm 0.07$), $P = 0.256$ | 0.04 ($\pm 0.06$), $P = 0.515$ | -0.02 ($\pm 0.02$), $P = 0.463$ |
| IFG | -0.02 ($\pm 0.07$), $P = 0.882$ | -0.13 ($\pm 0.06$), $P = 0.072$ | 0.03 ($\pm 0.02$), $P = 0.187$ |
| **Tap frequency variability (%): M5 response** |  |  |  |
| MI | -7.37 ($\pm 4.25$), $P = 0.256$ | -8.60 ($\pm 3.12$), **$P = 0.043$** | 2.30 ($\pm 1.12$), $P = 0.187$ |
| SMA | -8.74 ($\pm 3.87$), $P = 0.256$ | -8.60 ($\pm 2.99$), **$P = 0.043$** | 2.57 ($\pm 1.00$), $P = 0.148$ |
| MFG | -5.70 ($\pm 4.60$), $P = 0.350$ | -4.65 ($\pm 2.99$), $P = 0.201$ | 1.48 ($\pm 1.22$), $P = 0.423$ |
| IFG | 2.37 ($\pm 4.53$), $P = 0.804$ | 5.53 ($\pm 3.47$), $P = 0.201$ | -1.78 ($\pm 1.47$), $P = 0.423$ |
| **Tap frequency variability (%): M10 response** |  |  |  |
| MI | -8.09 ($\pm 4.81$), $P = 0.256$ | -10.75 ($\pm 4.97$), $P = 0.102$ | 2.89 ($\pm 1.62$), $P = 0.187$ |
| SMA | -1.63 ($\pm 5.57$), $P = 0.882$ | -3.49 ($\pm 4.69$), $P = 0.479$ | 0.47 ($\pm 1.84$), $P = 0.799$ |
| MFG | -9.56 ($\pm 4.70$), $P = 0.256$ | -6.72 ($\pm 4.35$), $P = 0.201$ | 2.64 ($\pm 1.47$), $P = 0.187$ |
| IFG | -0.40 ($\pm 4.65$), $P = 0.932$ | 6.26 ($\pm 3.63$), $P = 0.201$ | -0.73 ($\pm 1.35$), $P = 0.614$ |
| **Tap frequency variability (%): M20 response** |  |  |  |
| MI | -12.93 ($\pm 4.41$), $P = 0.105$ | -16.10 ($\pm 4.13$), **$P = 0.005$** | 4.60 ($\pm 1.37$), **$P = 0.028$** |
| SMA | -5.10 ($\pm 3.95$), $P = 0.343$ | -8.59 ($\pm 3.16$), **$P = 0.043$** | 1.88 ($\pm 1.03$), $P = 0.187$ |
| MFG | -7.07 ($\pm 4.14$), $P = 0.256$ | -7.07 ($\pm 3.27$), $P = 0.102$ | 2.08 ($\pm 1.08$), $P = 0.187$ |
| IFG | -1.45 ($\pm 4.30$), $P = 0.882$ | 5.29 ($\pm 3.21$), $P = 0.201$ | -0.80 ($\pm 1.03$), $P = 0.506$ |

Coefficient estimates in raw units, standard errors in brackets and adjusted P-values for the two predictors (stimulation amplitude and evoked response amplitude) as well as their interaction. Adjusted P-values <0.05 in bold. Regions of interest: MI = motor cortex, SMA = supplementary motor area, MFG = middle frontal gyrus, IFG = inferior frontal gyrus.

## Discussion

In this study we identified distinct cortical response patterns associated with different latencies. The M5 response involved motor cortex and supplementary motor area, while the M10 response was confined to the motor cortex and the M20 response again included the supplementary motor area, the middle and inferior frontal gyrus. Interestingly, the motor cortex M10 response was a predictor of finger tap frequency while the M5 and M20 related to a more regular movement profile indicating a fine-grained discrimination of movement by these responses.

So far, the cortical pattern of stimulation-evoked responses has only been studied to a limited extent, due to a lack of full cortical coverage of electrocorticography strips and a lack of spatial resolution of EEG recordings.^8,9,11,26^ Using MEG enabled us to localise the cortical responses evoked by subthalamic stimulation.

Earlier studies related responses between 2 and 10 ms to the activation of the hyperdirect pathway.^11,26^ These responses occur at three distinct latencies with a periodicity of about 2 ms.^7^ One possible explanation is that these responses are generated by a recurrent activation of layer V pyramidal neurons in the motor cortex following their antidromic activation.^27,28^ Our identified cortical responses at 5.6 and 11.8 ms support this hypothesis (Figure 1 C, D). Each of these responses involve the motor cortex and could therefore result from recurrent activation of motor cortex layer V neurons after antidromic cortical activation.

Moreover, a higher M5 response in the motor cortex and the supplementary motor area was indicative of a greater consistency in finger tapping and the motor cortex M10 response was predictive of a higher tap frequency. In animal models of Parkinson’s disease antidromic spiking of motor cortex layer V neurons as well as evoked motor cortex responses related to improved motor symptoms.^12,13^ Therefore, the M5 and M10 responses in our study could reflect the antidromic spiking (M5) and its after-effects (M10), relating to motor performance.

Stimulation-evoked cortical responses at longer latencies (> 20 ms) have been hypothesized to occur due to an orthodromic activation of the basal ganglia-thalamo-cortical loop.^11^ Our identified cortical responses at 21.8 ms appear within the inferior and middle frontal gyrus as well as the supplementary motor area. This is consistent with a polysynaptic activation involving various functional areas of the basal ganglia-thalamo-cortical loop.

It is important to note that only responses from motor cortex and supplementary motor area correlated with motor performance but even larger M20 responses in inferior and middle frontal areas did not (Figure 1 B, G). Thus, the stimulus-brain response relationship is restricted to the functionally relevant brain area of the task. Importantly, the patients did not perform any task during the MEG recording, but the relation to behavioural performance is based on a separate accelerometer recording with 130 Hz stimulation. Thus, these neural responses obtained in a task-free manner could be used as markers for optimal stimulation settings.

One limitation of MEG studies with DBS patients is the change from clinically-used monopolar to bipolar DBS to reduce stimulation related artefacts.^16,26^ To improve comparability with clinical settings, we used monopolar stimulation. However, cortical responses of less than 3 ms might still be contaminated by the monopolar stimulation artefact. Therefore, we did not include these responses in our tested models. Another limitation of our study is the exclusive focus on finger tapping as the behavioural marker of motor performance.

## Conclusion

STN stimulation-evoked cortical responses could inform clinical programming in a relatively precise manner, because they relate to different movement profile characteristics, i.e. movement velocity and regularity. The amplitude of the motor cortex response could be especially useful to select a clinically effective stimulation contact. Due to the low stimulation frequency, the recording does not induce side effects. Prospective clinical studies are needed to validate the applicability of stimulation-evoked responses to aid DBS programming.

## Abbreviations

DBS: Deep Brain Stimulation
ECoG: Electrocorticography
MEG: Magnetoencephalography
UPDRS: Unified Parkinson’s disease Rating Scale
STN: Subthalamic Nucleus
tSSS: temporal signal space separation

## Acknowledgements

We thank Pia Hartmann and Luisa Spallek for their assistance during recordings and Johannes Pfeifer for critically reviewing the manuscript. We thank all of the patients for their participation.

